# *C9orf72*-derived poly-GA DPRs undergo endocytic uptake in iNPC-derived astrocytes and spread to motor neurons

**DOI:** 10.1101/2021.10.11.463891

**Authors:** Paolo M. Marchi, Lara Marrone, Laurent Brasseur, Luc Bousset, Christopher P. Webster, Marco Destro, Emma F. Smith, Christa G. Walther, Victor Alfred, Raffaele Marroccella, Darren Robinson, Allan C. Shaw, Lai Mei Wan, Andrew J. Grierson, Stephen J. Ebbens, Kurt J. De Vos, Guillaume M. Hautbergue, Laura Ferraiuolo, Ronald Melki, Mimoun Azzouz

## Abstract

Dipeptide repeat proteins (DPRs) are aggregation-prone polypeptides encoded by the pathogenic G4C2 repeat expansion in the C9orf72 gene, the most common genetic cause of amyotrophic lateral sclerosis and frontotemporal dementia (ALS/FTD). In this study, we focus on the role of poly-GA DPRs in disease spread. We demonstrate that recombinant poly-GA oligomers can directly convert into solid-like aggregates and form characteristic &[beta]-sheet fibrils in vitro. To dissect the process of cell-to-cell DPR transmission, we closely follow the fate of poly-GA DPRs in either their oligomeric or fibrillized form after administration in the cell culture medium. We observe that poly-GA DPRs are taken up via dynamin-dependent and - independent endocytosis, eventually converging at the lysosomal compartment and leading to axonal swellings in neurons. We then use a co-culture system to demonstrate astrocyte-to-motor neuron DPR propagation, showing that astrocytes may internalise and release aberrant peptides in disease pathogenesis. Overall, our results shed light on the mechanisms of poly-GA cellular uptake and cell-to-cell propagation, suggesting lysosomal impairment as a possible feature underlying the cellular pathogenicity of these DPR species.

## Introduction

The polymorphic hexanucleotide repeat expansion (HRE) in the *C9orf72* gene is the major genetic cause of amyotrophic lateral sclerosis/frontotemporal dementia (ALS/FTD) (Renton *et al*, 2011; DeJesus-Hernandez *et al*, 2011). The pathogenic HRE consists of hundreds to thousands of GGGGCC (G4C2) repeats, located in the first intron of the *C9orf72* gene (van Blitterswijk *et al*, 2013). A crucial driver of *C9orf72*-mediated ALS/FTD pathology is the unconventional repeat-associated non-AUG (RAN) translation of the HRE into five toxic dipeptide repeat (DPR) species. *Post-mortem* tissue of ALS/FTD patients contains ubiquitin- and p62-positive DPR inclusions predominantly in the frontal cortex, hippocampus and cerebellum of neuronal cells (Mori *et al*, 2013; Ash *et al*, 2013; Schludi *et al*, 2015; Saberi *et al*, 2018), with rare occurrence in glia (Rostalski et al., 2019).

Among the five different DPRs generated by RAN translation, poly-GA appears to be the most abundantly detected as well as one of the most toxic DPR species (Zhang et al., 2014; Mackenzie et al., 2015). Poly-GA toxicity has been documented both *in vitro* and *in vivo* (Lee et al., 2017; Ohki et al., 2017; Nihei et al., 2019) and correlates with motor deficits, cognitive defects and inflammatory response in mice (Zhang et al., 2016; Schludi et al., 2017; LaClair et al., 2020). Initial efforts to identify molecular mechanisms of poly-GA toxicity revealed that these DPR species interact with components of the Ubiquitin-Proteasome System, such as p62, ubiquilin-1, ubiquilin-2, HR23 (May et al., 2014; Schludi et al., 2015; Zhang et al., 2016) and specifically lead the 26S proteasome to stalled degradation (Guo et al., 2018). Emerging evidence shows that Poly-GAs can also rapidly spread throughout the *Drosophila* brain in a repeat length- and age-dependent manner (Morón-Oset *et al*, 2019), in agreement with the ability of Poly-GAs to spread and drive cytoplasmic mislocalization and aggregation of TDP- 43 in cell cultures (Chang *et al*, 2016; Westergard *et al*, 2016; Zhou *et al*, 2017; Khosravi *et al*, 2020).

In this work, we used an *E. Coli*-based system for the production of poly-GA oligomers of 34 repeats. We first illustrate the process of poly-GA oligomer coalescence into solid-like species. Importantly, poly-GA oligomers show a distinct ability to form fibrillar species of amyloid nature with characteristic β-sheets *in vitro.* We next explored whether different poly-GA species (poly- GA oligomers vs poly-GA fibrils) could originate distinct features in terms of cellular uptake and cell-to-cell propagation. We show that poly-GA oligomers (and not fibrillary GA) access cells despite dynamin inhibition and escape lysosomal degradation.

Upon active uptake, both poly-GA oligomers and fibrils are transported to lysosomes, which may become aberrantly enlarged and static, leading to axonal swellings in neurons. When astrocytes and motor neurons are co-cultured, DPR species are promptly internalized by astrocytes and spread to neuronal units. Our data highlights the steps of poly-GA DPR spread *in vitro*, suggesting lysosomal impairment as a potential pathogenic mechanism underlying disease.

## Results

### Recombinant poly-GA aggregation into oligomeric and fibrillar assemblies

To study the potential role of poly-GA DPRs in ALS/FTD spread, we first set up an expression system in *E. coli* for the production of recombinant poly-GA DPRs labelled with Atto-550 or Atto-647N dyes (see Materials & Methods). Additionally, we included poly-PA DPRs for comparison. GA/PA-repeat immunoreactivity and fluorescent labelling were confirmed by resolving the generated samples by protein gel electrophoresis or dot-blot **(Fig S1A-C)**.

We first investigated the aggregation of poly-GA and -PA at different concentration using confocal microscopy. Both ATTO550-labeled DPRs coalesced into microscopic protein clusters under low-salt concentrations and without the addition of any molecular crowders **(Figs 1A and B)**. Clusters formation was much faster for poly-PA than poly-GA, and grew with increasing protein concentration (1µM, 10µM, 20µM) **(Figs 1Ai and Bi)** and incubation length (0h, 2h, 24h) for both DPRs **(Figs 1Aii and Bii)**. The generated poly-GA and -PA clusters differed significantly in their shape and size, hence we used Z-stack confocal microscopy and CMLE deconvolution to view their 3D-volume and -surface rendering **(Figs 1Aiii and Biii)**. The 3D-reconstructed poly-GAs showed an irregular and compact solid-like structure **(Movie S1)**. In sharp contrast, the 3D-reconstructed poly-PAs were made of very small spherical particles (circularity = 0.95; **Fig S1D**) resembling liquid-like droplets **(Movie S2)**.

**Figure 1.**
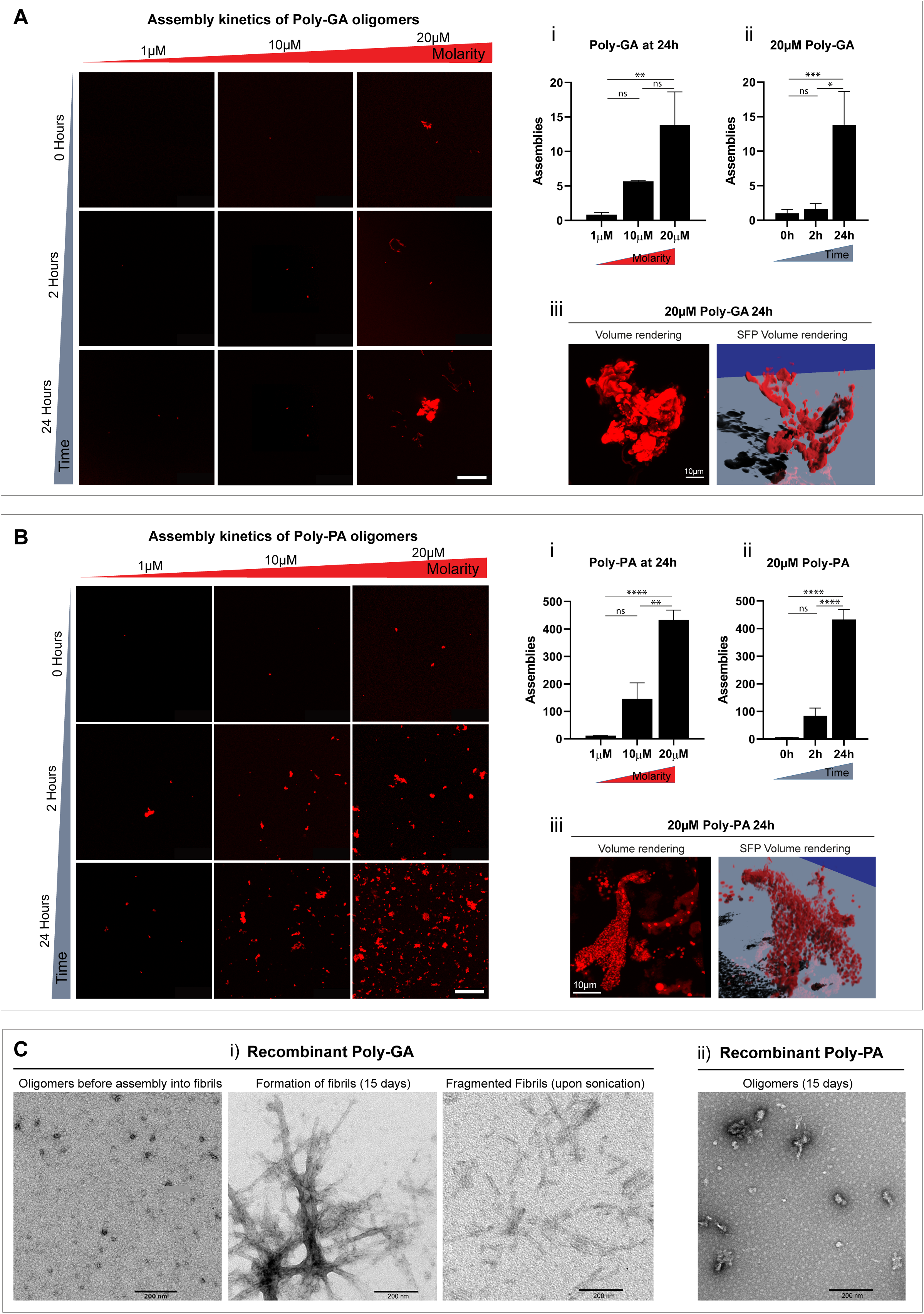
Poly-GA oligomers form solid-like structures and assemble into characteristic ß-sheet fibrils. **A)** Aggregation of poly-GA oligomers *in vitro*, with relative quantification of the number of poly-GA assemblies formed upon increasing molarity (i) or time (ii). The largest structures formed at 20µM 24h are shown in 3D rendered images (iii); corresponding movie is shown in **Movie S1**. **B)** Aggregation of poly-PA oligomers *in vitro*, with relative quantification upon increasing molarity (i) or time (ii). The largest structures formed at 20µM 24h are shown in 3D rendered images (iii); corresponding movie is shown in **Movie S2**. For both A) and B): Scale bar = 50 μm. Bar graphs of mean ± SEM. One-way ANOVA with Tukey’s multiple comparisons test. *P ≤ 0.05, **P ≤ 0.01, and ****P ≤ 0.0001; N=2. **C)** Electron micrographs showing that poly-GA oligomers form characteristic fibrils after 15 days *in vitro* (i), unlike poly-PA oligomers (ii). Poly-GA fibrils are shown before and after fragmentation, which drives the production of fragmented fibrils. Scale bar = 200 nm.

Using longer incubation times, we then aimed to investigate whether Poly-GA oligomers could grow *in vitro* into β-sheet fibrils. As observed by transmission electron microscopy, we noticed that, while poly-PA formed non-fibrillary amorphous assemblies, poly-GA assembled into fibrillar structures within 15 days of incubation **(Fig 1C)**. Notably, the poly-GA fibrillar DPRs used in this study were extensively characterized for their amyloid β-sheet content by Fourier-transform infrared (FTIR) spectroscopy (Brasseur et al., 2019). In summary, our poly-GA oligomers produced distinctive solid-like assemblies, nucleation growth and 3D-architecture and uniquely assembled into characteristic β-sheet fibrils.

### Oligomeric and fibrillar poly-GA DPRs use distinct entry-routes in glia

To evaluate DPR uptake, we exposed various cell types (HEK293T, HeLa, iNPC-derived human astrocytes, human fibroblasts) to our recombinant DPRs for 1h, 2h and 4h. Using high-throughput confocal microscopy, we then quantified the number of largely visible DPR aggregates in these cell types. Our results on ∼8,000 cells/culture showed that in all these cell lines poly-GA oligomers are taken up more readily than poly-GA fibrils and poly-PA oligomers. Additionally, for all DPR species and across all cell lines, the uptake increases with time **(Fig 2A)**.

**Figure 2.**
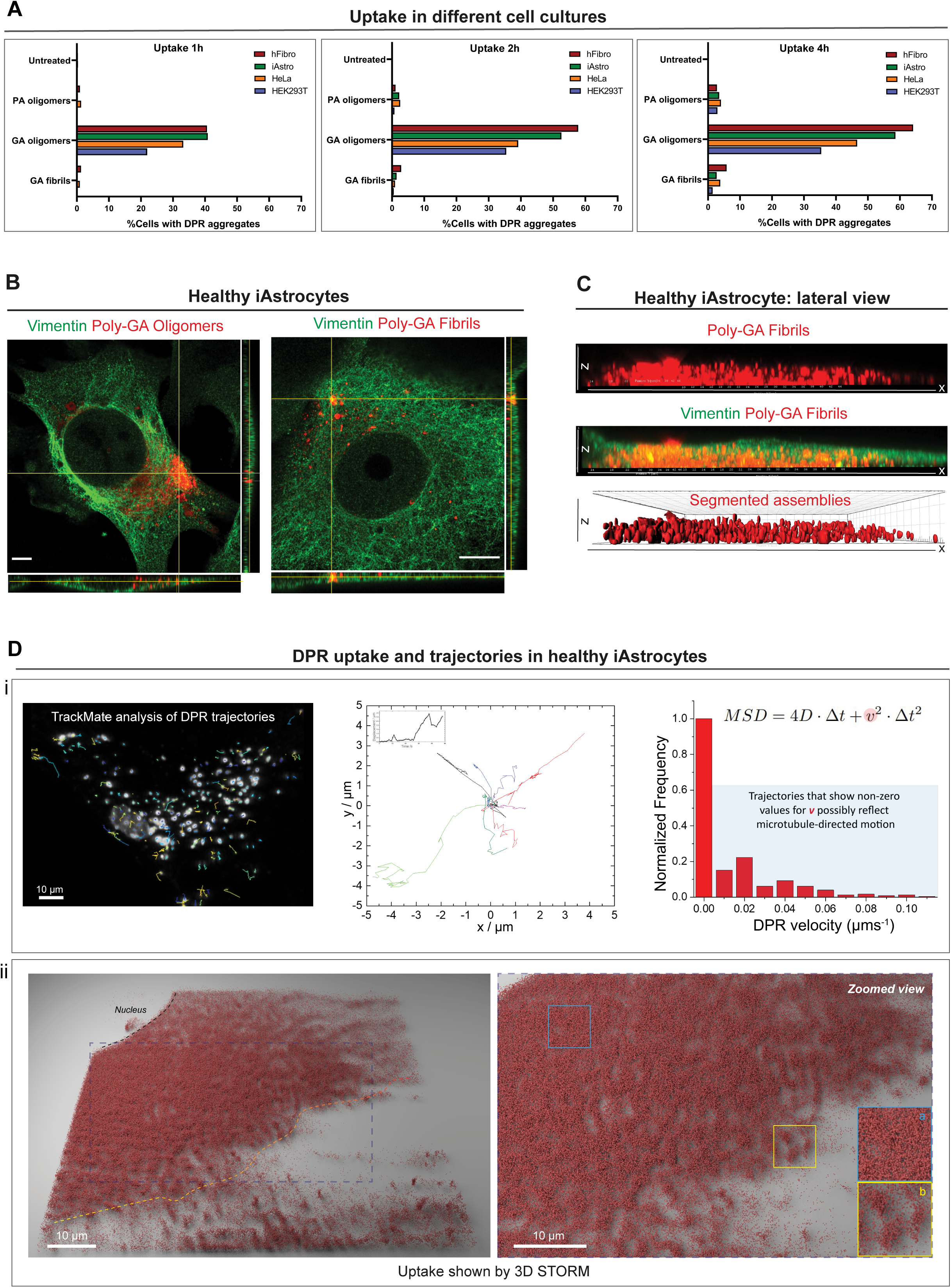
DPR uptake in various cell lines and in healthy iAstrocytes. **A)** Quantitative analysis of the %cells displaying large DPR aggregates overtime (1h, 2h, 4h). Different cell cultures were used in this experiment such as human fibroblasts *(red)*, iAstrocytes *(green)*, HeLa cells *(orange)*, and HEK293T cells *(blue)*. ∼8’000 cells/culture were analysed; N=3. **B)** Orthogonal views from airyscan microscopy show the uptake of poly-GA oligomers and poly-GA fibrils in vimentin-stained healthy iAstrocytes after 24h exposure. Scale bar = 10μm. **C)** 3D-rendered lateral view of a single vimentin-stained healthy iAstrocyte shows the uptake of poly-GA fibrils through the xz dimension. Corresponding movies are shown in **Movie S3 and S4**. **D)** Mean Square Displacement analysis on DPR trajectories after live-imaging in healthy iAstrocytes (i). Some tracks, analysed with TrackMate, are shown in the x,y space; corresponding movie shown in **Movie S5**. The quantification of DPR average velocity *v* is displayed in the red graph (i). 3D-STORM imaging was used to visualize ATTO647N-DPRs in healthy iAstrocytes after 24h uptake (ii). The zoomed view shows how DPRs can be densely clustered *(inset a)* but also compartmentalized in spherical structures *(inset b)*.

Because increasing evidence implicates astrocytes as significant non-cell autonomous contributors of *C9orf72*-associated ALS/FTD pathogenesis (Varcianna *et al*, 2019), we aimed at better investigating poly-GA DPR uptake in healthy iNPC-derived human astrocytes (herein referred to as iAstrocytes). We firstly confirmed DPR internalization in iAstrocytes by using Z-stack airyscan confocal microscopy, orthogonal views and 3D-volume rendering **(Figs 2B and C; Movies EV3 and EV4)**. Uptake was further evidenced by live-cell imaging of iAstrocytes and subsequent quantification of DPR average velocity (*v)* with a Mean Square Displacement analysis (see Materials & Methods_*MSD(Δt) analysis*). Our analysis showed non-zero values for *v*, which indicates that DPR motion is not solely due to random diffusion, but an active force is also contributing **(Fig 2Di)**. This force resembles microtubule-mediated motion **(Movie S5)**, as also suggested by the fact that the microtubule de-polymerising agent nocodazole induces cellular relocation of these DPRs **(Fig S2A)**. We further showed DPR uptake in glia by the use of 3D-STORM imaging **(Fig 2Dii)** and trypan blue quenching **(Fig S2B)**, and DPR binding to cells by flow cytometry **(Fig S2C)** and anti-V5 dot-blot **(Fig S2D)**.

To test the potential involvement of endocytosis in DPR uptake, we performed a generalised block of all endocytic pathways by lowering the culture temperature to 4°C prior to fixation (Harding *et al*, 1983). To confirm the successful inhibition of endocytosis we visually monitored the uptake of transferrin, a well-established marker of the coated pit pathway (Ehrlich *et al*, 2004; Hanover *et al*, 1984). Interestingly, low-temperatures reduced the uptake of poly-GA oligomers by 2.2-fold and of poly-GA fibrils by ∼8-fold (P**** ≤ 0.0001) in iAstrocytes **(Fig 3A)**. We next inhibited dynamin-dependent endocytosis by the use of dynasore (Macia *et al*, 2006; Kirchhausen *et al*, 2008), and this resulted in a 2.4-fold uptake reduction of poly-GA fibrils (P**** ≤ 0.0001) but no change in poly-GA oligomers uptake **(Fig 3B)**. Upon comparison with other DPR oligomeric species (such as poly-PA), we observed that specifically poly-GA oligomers do not use dynamin-dependent endocytosis for cell entry. To better investigate the differences in uptake between oligomeric vs fibrillar poly-GA DPRs, we evaluated the accumulation of these proteins in endolysosomal organelles following 24h from administration. We exposed healthy iAstrocytes to poly-GA fibrils or oligomers (0.5µM for 24h) and quantified the degree of colocalization with LAMP1-stained endolysosomes by dual-colour dSTORM with a lateral resolution of 50 nm for both channels **(Fig 3C and Fig S3A-C)**. The analysis revealed that while ∼17% of the input poly-GA fibrils co-localized with endolysosomes, only less than 5% poly-GA oligomers did (3.4-fold difference; P*** ≤ 0.001). Additionally, LAMP1 organelles showed higher enrichment for poly-GA fibrils than for poly-GA oligomers (1.6-fold difference; P*** ≤ 0.001) **(Fig 3C)**. Together, these findings suggest that DPR uptake is present in various cell culture systems and endocytosis plays a role in DPR uptake in iAstrocytes.

**Figure 3.**
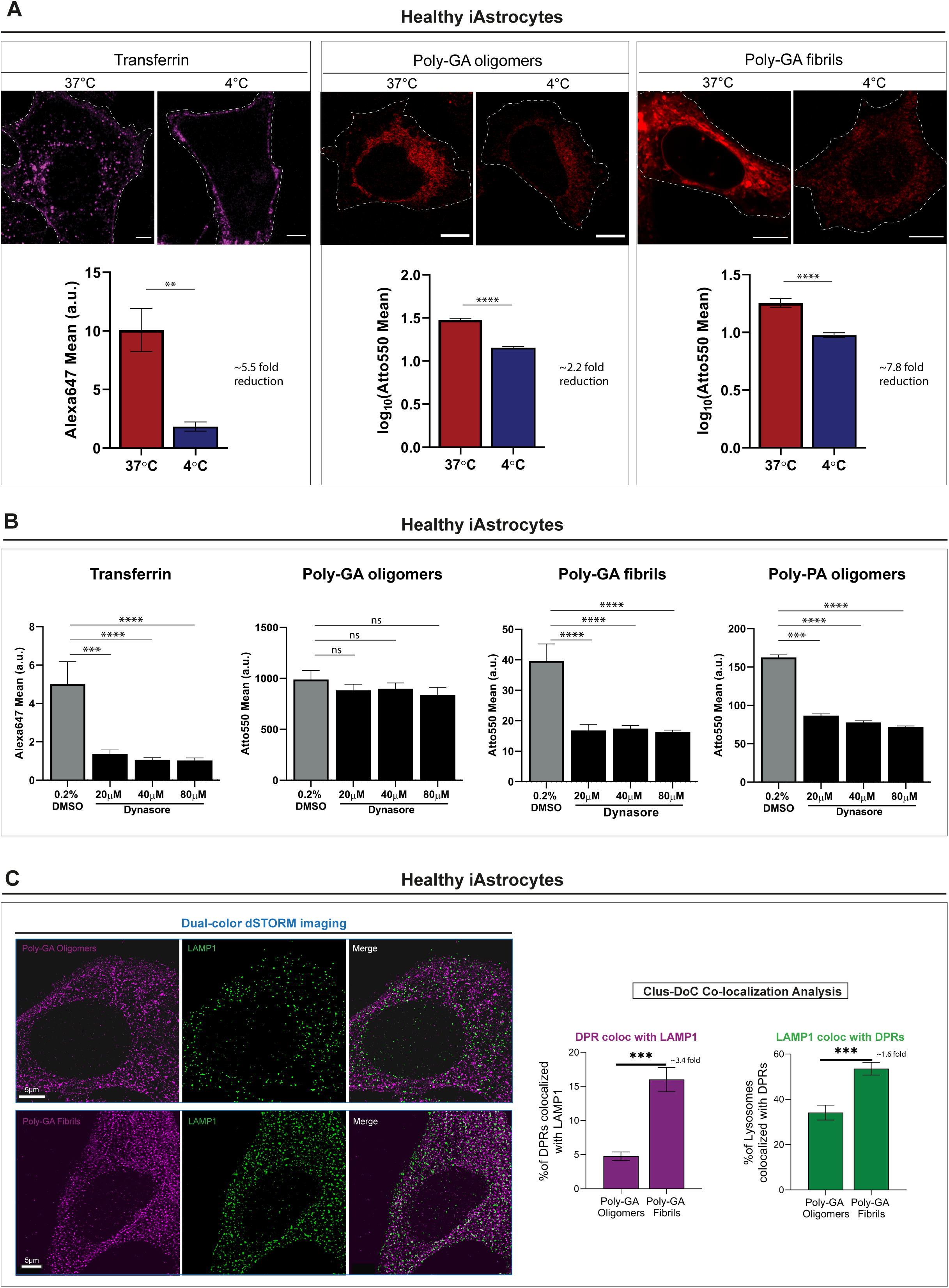
Oligomeric vs fibrillar poly-GA DPRs entry-routes in glia. **A)** Confocal images of Alexa647-Transferrin (control) and ATTO550-Poly-GA aggregates after 1h uptake at 37°C or at 4°C in healthy iAstrocytes. Quantification of the log_10_ transformed- Mean grey values is reported in bar graphs of mean ± SEM. ≥300 cells/condition; N=4. Kolmogorov-Smirnov non-parametric test after testing normal distribution with the Shapiro- Wilk test. **P ≤ 0.01, and ****P ≤ 0.0001. **B)** Quantification of the Mean grey values of Alexa647-Transferrin (control) and ATTO550-DPRs upon 1h treatment with Dynasore (or 0.2% DMSO). Bar graphs of mean ± SEM. ≥250 cells/condition; N=3. Kruskal-Wallis non- parametric test with Dunn’s multiple comparisons after testing normal distribution with the Shapiro-Wilk test. ***P ≤ 0.001, and ****P ≤ 0.0001. **C)** Healthy iAstrocytes were imaged by dual-colour STORM after 24h incubation with ATTO647N-PolyGA oligomers or fibrils *(magenta)* and anti-LAMP1 staining *(green)*. Clus-DoC co-localization analysis for Poly-GA relative to LAMP1 *(magenta graph)*, and LAMP1 relative to Poly-GA *(green graph)* shows the respective %colocalized molecules (among total molecules detected). Scale bar = 5 μm. Bar graphs of mean ± SEM; graphs are indicative of 30 ROIs (4μm x 4μm) per condition chosen only in artefact-free regions. Unpaired two-tailed t-test with Welch’s correction. ***P ≤ 0.001; N=3.

### Poly-GA DPRs accumulate into enlarged lysosomes and into axonal swellings in neurons

Our DPR-endolysosomal colocalization finding led us to hypothesize that poly-GA oligomers might have the ability to escape endolysosomal routes and trigger cellular toxicity via lysosomal disruption. Therefore, we next sought to test whether poly-GAs could cause lysosomal impairment in cortical neurons, which are cells that undergo degeneration in ALS-FTD and display a large number of DPR inclusions in *post-mortem* tissue of ALS-FTD patients (Mackenzie et al., 2015).

After growing primary mouse cortical neurons into a microfluidic chamber system, we used live-imaging and detected poly-GA uptake in the soma (point of DPR exposure) as well as in the proximal axon. Poly-GA DPRs shuttle along axons by a mixture of Brownian motion, directed motion and constrained motion **(Fig S4A).** Constrained motion was the predominant DPR motion throughout the axons; suggesting that these proteins are mostly anchored to static axonal structures. We indeed found that poly-GA DPRs accumulated in large axonal swellings, implying a failure of their axonal transport **(Fig 4Ai)**. Interestingly, by zooming into few axonal swellings with higher resolution, we observed the presence of small poly-GA proteins erratically moving within each axonal swelling **(Fig 4Aii**, **Movie S6).** We then used the lysosomal dye LysoTracker Green and observed that some of these axonal swellings contain Poly-GA DPRs colocalizing with lysosomal organelles **(Fig S4B**, **Movie S7)**. We next performed a thorough co-localization analysis between all the DPRs and the lysosomes contained in the proximal axons. By using color deconvolution algorithms, we could separately analyse two lysosomal populations: one which does not co-localize with poly-GA DPRs (“Non-Colocalized Lysosomes”, NCLs) and one showing co-localization (“Colocalized Lysosomes”, CLs) **(Movie S8)**. Interestingly, CLs displayed reduced displacement, reduced speed, and increased size compared to NCLs in proximal axonal regions (**Fig 4B**).

**Figure 4.**
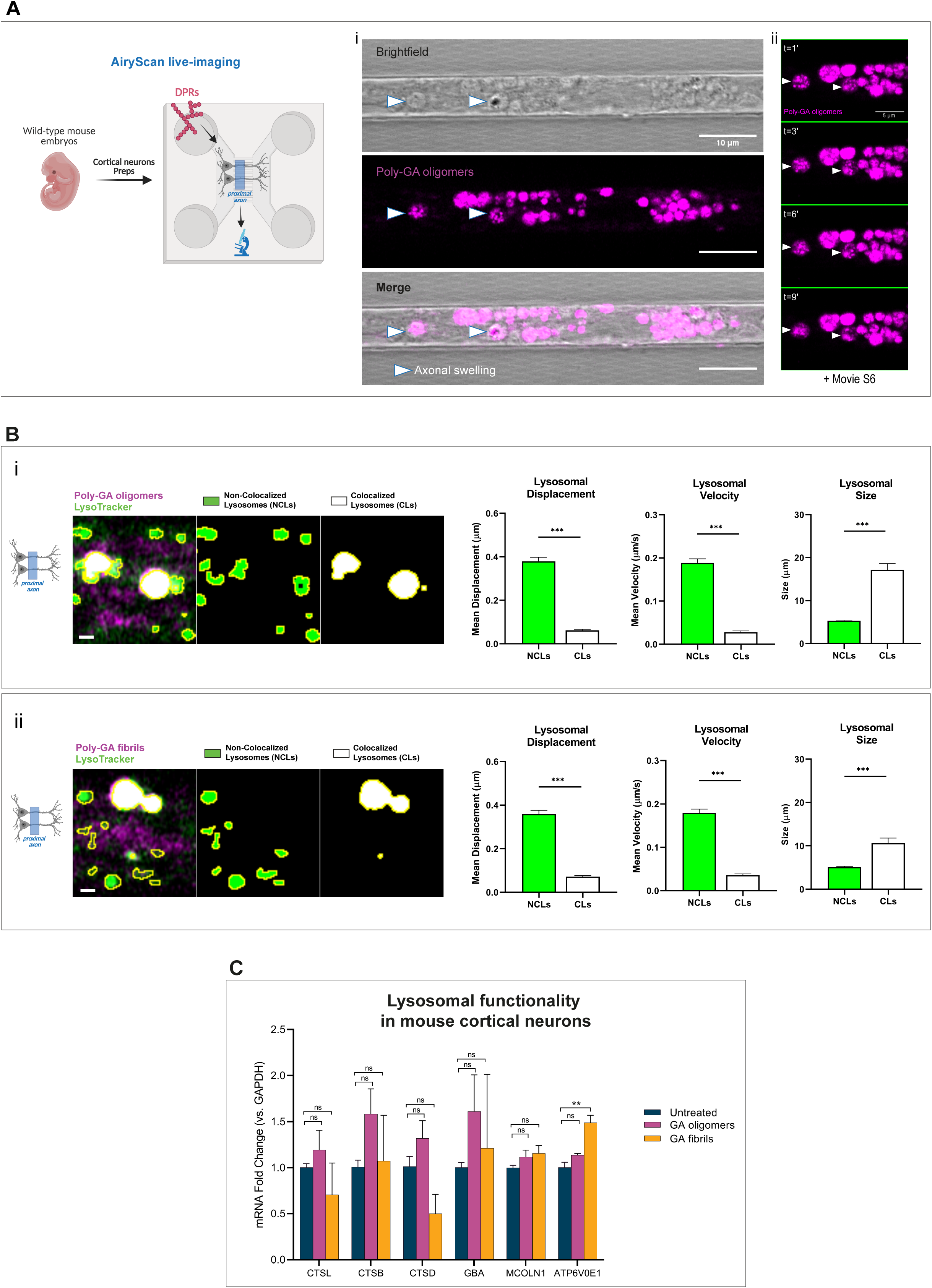
Poly-GA aggregates colocalize with endolysosomal organelles after uptake and disrupt lysosomal homeostasis in neurons. **A)** Confocal images showing accumulation of poly-GA assemblies in large axonal swellings *(arrow heads)* along the axons of primary mouse cortical neurons. A combination of brightfield imaging and fluorescence detection of the ATTO550-labeled Poly-GA assemblies was used to detect these swellings specifically in the axons residing in the microfluidic chamber microgrooves (i). By zooming into few axonal swellings with higher resolution (airyscan mode), during live-imaging, we report the presence of small poly-GA assemblies particles erratically moving within each axonal swelling overtime (ii); corresponding movie is shown in **Movie S6**. Figure of Cortical neurons Preps created with BioRender.com under academic license. **B)** Co-localization analysis between all the DPRs and the lysosomes contained in the cortical neurons’ proximal axons. By color deconvolution we separated “Non-Colocalized Lysosomes” (NCLs) and “Colocalized Lysosomes” (CLs) to analyse displacement, speed and size. Scale bars = 1 μm. Bar graphs of mean ± SEM. Unpaired two-tailed t-test with Welch’s correction. ***P ≤ 0.001; N=3. Figure of Proximal axon created with BioRender.com under academic license. **C)** Quantification of relative expression levels after qRT-PCR for genes involved in lysosomal metabolism. RNA was isolated from primary mouse cortical neurons (10 days in vitro) exposed or not to poly-GA oligomers or poly-GA fibrils for 24h, retrotranscribed and subjected to quantitative real-time PCR. Only the transcriptional levels of ATP6V0E1 (a component of the V-ATPase) were increased upon treatment with poly-GA fibrils compared to control untreated neurons. One-way ANOVA with Tukey’s multiple comparisons test. **P ≤ 0.01; N=3.

To further investigate signs of lysosomal-induced damage and dissect potential differences between poly-GA fibrils and poly-GA oligomers, we exposed primary cortical neurons to these DPR species for 24h, followed by RNA extraction and qRT-PCR for genes involved in lysosomal metabolism. While transcripts encoding various cathepsins (CTSL, CTSB, CTSD), a lysosomal hydrolase (GBA), and a cation-permeable lysosomal channel (MCOLN1) did not show variation upon DPR exposure, the transcriptional levels of ATP6V0E1 (a component of the V-ATPase) were increased upon treatment with poly-GA fibrils **(Fig 4C)**. To sum up, our findings suggest that poly-GA DPRs disrupt lysosomal homeostasis in the axons of cortical neurons.

### Poly-GA DPRs undergo astrocyte-to-motor neuron spread

Despite neuron-to-astrocyte transmission has been documented (Westergard *et al*, 2016), the role of glial cells in DPR propagation remains mostly unexplored, and no study to our best knowledge has yet unveiled whether DPRs can propagate from astrocytes to neurons. To test the possibility of this directionality, we established a co-culture system between iAstrocytes and Hb9-GFP mouse motor neurons. Briefly, we treated healthy iAstrocytes with 1 µM DPRs for 24h, performed a number of PBS washes to remove remaining assemblies, and subsequently plated Hb9-GFP mouse motor neurons over the astrocyte layer. We kept this co-culture system for 48h before fixation and confocal imaging **(Fig 5A)**. We observed that, after astrocytic uptake, DPR assemblies underwent astrocyte-to-neuron propagation as confirmed by orthogonal views **(Fig 5A).** Different DPRs showed different percentages of propagation to motor neurons, with poly-GA fibrils being the most efficient at spreading (detected in 24% of motor neurons) when compared to oligomeric species (poly-GA: 4%; poly-PA: 7%) **(Fig 5Bi)**. We then verified the release of DPRs into the conditioned medium (CM) by spectrophotometrically quantifying ATTO550 fluorescence levels **(Fig 5Biii)**. Eventually, we investigated whether the astrocyte-to-neuron transmission of DPRs could contribute to motor neuron damage as a non-cell autonomous effect. However, no evident cytotoxicity was found in motor neurons upon co-cultures with iAstrocytes that contained and transmitted DPRs **(Fig 5Biii)**.

**Figure 5.**
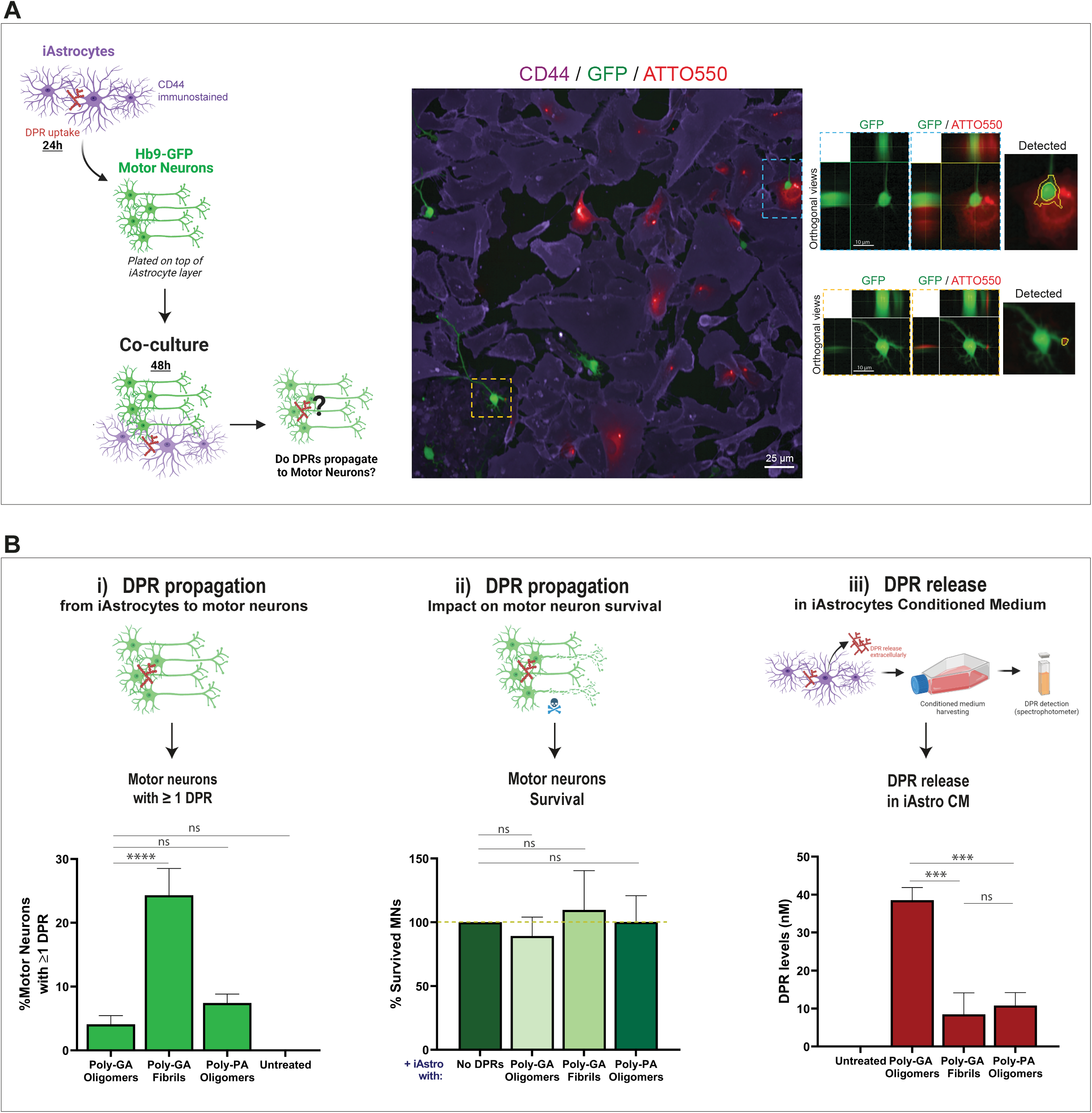
Alanine-rich DPRs undergo astrocyte-to-motor neuron propagation. **A)** Schematic representation of the iAstrocytes-MNs co-culture system *(left panel)*. Orthogonal views from confocal images show the presence of ATTO550 DPRs in the GFP-positive MNs; CD44 is shown in violet as the astrocytic marker *(middle and right panels)*. Figure of cells created with BioRender.com under academic license. **B)** Quantification of the %motor neurons containing at least one DPR aggregate in the various 48h co-culture systems (i). Quantification of motor neuron survival upon 48h co-culture with iAstrocytes containing and transmitting DPRs (ii). For both (i) and (ii): bar graphs of mean ± SEM. One-way ANOVA with Tukey’s multiple comparisons test. ∼120 neurons/condition. ns = P > 0.05; N=3. Quantification of DPR levels present in the conditioned medium of healthy iAstrocytes, via spectrophotometric analysis (iii). One-way ANOVA with Tukey’s multiple comparisons test. ns = P > 0.05, and ***P ≤ 0.001; N=4. Figure of cells created with BioRender.com under academic license.

Taken together, our results show that iAstrocytes can efficiently release alanine-rich DPRs into the culture medium and, when co-cultured with motor neurons, they are able to transmit DPRs to neuronal cells. Although we did not detect any evident cell death following transmission with our system, we speculate that astrocytes may act as hubs for the internalisation and release of aberrant peptides in ALS.

## Discussion

Pathological protein aggregates in neurons and glia are a hallmark of most neurodegenerative diseases (Eisele *et al*, 2015). In the context of *C9orf72*-ALS/FTD, aggregation of the RAN translation-derived Poly-GA DPRs is one of the proposed mechanisms for inducing proteasome impairment, DNA damage, cognitive disability, motor deficits, pro-inflammatory responses and neurodegeneration, as shown by numerous *in vitro* and *in vivo* studies (Mori *et al*, 2013; Zhang et al., 2016; Schludi et al., 2017; Guo et al., 2018; Nihei et al., 2019; LaClair et al., 2020). In proteinopathies, the initial stages of the aggregation process involve the nucleation of heterogeneous oligomeric species, which are then thought to progressively grow into compact cross-β fibrils (Serio et al., 2000; Bernstein et al., 2009; Narayan, et al., 2012; Cohen et al., 2013). Increasing evidence suggests that oligomeric species, and not fibrils, would act as the major pathogenic culprit in neurodegenerative disease, with their seeding activity being particularly powerful in the earliest stages of pathogenesis (Walsh et al., 2002; Campioni et al., 2010; Winner et al., 2011; Cremades et al., 2012; Ye et al., 2017; Uhlmann et al., 2020).

In our work, we began our analysis of poly-GA DPR spread by investigating the process of poly-GA oligomerization and fibril formation. We firstly showed that the *in vitro* coalescence of poly-GA and poly-PA oligomers increased in correlation with protein concentration and incubation length. However, while poly-GAs formed large and solid-like assemblies, poly-PAs produced small spherical liquid-like droplets. This result is in agreement with the reduced toxicity of poly-PA DPR species (Mizielinska et al, 2014; Vanneste et al., 2019), since maintenance of liquid-phase homeostasis was proposed to be non-pathogenic in protein aggregation (Patel *et al*, 2015; Peskett *et al*, 2018; Ray *et al*, 2020). Next, we aimed to investigate whether Poly-GA oligomers could form β-sheet fibrils *in vitro*. Our results showed that poly-GA readily assembled into fibrillar structures within 15 days of incubation, as opposed to poly-PA DPRs. Since the transition to β-sheet fibrils exposes hydrophobic amino acid residues (Landreh *et al*., 2016), we speculate that poly-GA could lead to significant problems of insolubility as previously described in cell culture studies (Zhang et al., 2014; Lee et al., 2017; Ohki et al., 2017; Nihei et al., 2019; LaClair et al., 2020).

We next wanted to perform comparative experiments featuring recombinant poly-GA oligomers and fibrils in order to explore whether differences in the aggregation stage could produce differences in cellular uptake, toxicity and cell-to-cell propagation. We first exposed various cells to poly-GA DPRs using a concentration range of 0.5-1µM, which presumably recapitulates physiologically relevant conditions (Meeter *et al*, 2018). Interestingly, we showed that poly-GA oligomers are taken up more easily than poly-GA fibrils or other oligomeric species such as poly-PA. We next demonstrated that at least a fraction of poly-GA DPRs is internalized by endocytosis in iAstrocytes: low-temperature exposure (which inhibits all endocytic pathways) reduced the uptake of poly-GA oligomers by 2.2-fold and of poly-GA fibrils by ∼8-fold. Intriguingly, when more specific pathways of endocytosis were pharmacologically inhibited, such as those depending on the GTPase protein dynamin, only the uptake of poly-GA fibrils was reduced but no change was observed in poly-GA oligomers uptake. This result might be explained by an intrinsic capability of small oligomers to escape endolysosomal clearance, perhaps by triggering lysosomal permeabilization and cellular toxicity (Lee et al, 2008; Tomic et al, 2009; Kandimalla et al, 2009). To test this hypothesis, we used super-resolution imaging to investigate the degree of co-localization between poly-GA DPRs and LAMP1-stained endolysosomal organelles in iAstrocytes and indeed found reduced lysosomal co-localization of poly-GA oligomers compared to poly-GA fibrils.

To explore potential lysosomal-induced damage driven by poly-GA oligomers vs poly-GA fibrils, we chose to incubate these DPR species with mouse cortical neurons. When administered to the neuronal soma in a microfluidics system, these DPRs were taken up and transported along the axons, accumulating in stalling lysosomes leading to axonal swellings. Importantly, the lysosomal population which co-localized with DPRs presented abnormalities in motility, speed and size compared to the non-colocalized counterpart. The regulation of lysosomal motility and size is particularly crucial in neurons because of their polarity: lysosomes need to access specific cytoplasmic locations to perform their various functions and a correct lysosomal size is crucial for fission and fusion events. Mutations in components regulating lysosomal motility cause various neurological disorders (Pu *et al*., 2016), and the size of lysosomes is altered in several human diseases (de Araujo *et al*., 2020). Our results showing that poly-GA DPRs accumulate in aberrantly immotile and enlarged lysosomes suggests that these proteins might impair the function of these organelles either directly or, alternatively, indirectly by disrupting key lysosomal partners. For the latter hypothesis, it was recently shown that arginine-rich DPRs impede translocation of dynein and kinesin-1 motor complexes and bind microtubules promoting their pausing and detachment (Fumagalli *et al*., 2021). We speculate that similar mechanisms could be behind the behavior observed following poly-GA DPR internalization, as our data showed that poly-GAs are cellularly relocated upon nocodazole treatment, thus suggesting the presence of polyGA- microtubule interactions. Our data on lysosomal functionality also showed that the transcript levels of ATP6V0E1 were increased upon 24h exposure of poly-GA fibrils compared to untreated or poly-GA oligomers exposure. ATP6V0E1 is a component of the multi-subunit ATPase enzyme, which is crucial to promote the acidification of intracellular organelles in eukaryotes. Upregulating ATP6V0E1 may constitute a cellular response to facilitate the clearance of poly-GA fibrils, potentially by maintaining or enhancing lysosomal pH acidic gradients via V-ATPase activity.

We finally sought to investigate whether poly-GA oligomers and poly-GA fibrils could show differences in the ability to undergo cell-to-cell propagation, a process previously observed for various DPR species in neuronal cell cultures (Chang *et al*, 2016; Westergard *et al*, 2016; Zhou *et al*, 2017; Khosravi *et al*, 2020) as well as in the Drosophila nervous system (Morón- Oset *et al*, 2019). We proceeded to explore these questions by establishing a co-culture system between iAstrocytes and Hb9-GFP mouse motor neurons; this system was used because of two main reasons: (i) the largely unexplored role of glia in DPR propagation with relation to neurons, (ii) and the previously described role of astrocytes as “hubs” for intercellular propagation of protein aggregates (Victoria et al, 2016; Loria et al, 2017; Wang et al, 2019). Our results showed that poly-GA DPRs undergo astrocyte-to-neuron propagation,

with the fibrils being 6-times more efficient in transferring to motor neurons compared to oligomers. Thus, DPRs at later stages of aggregation might be more prone to transfer from affected to naïve cells, which is in agreement with previous findings related to other amyloid proteins (Rey et al., 2019). Somewhat surprisingly, we were not able to detect any cytotoxicity in co-cultured motor neurons associated to poly-GA DPRs. Cytotoxicity might perhaps require the synergistic presence of other DPR species (such as poly-GR and poly-PR), which were not included in our experimental settings. Nonetheless, our data propose an important role for astrocytes in the transmission of *C9orf72*-derived DPRs to neighbouring motor neurons. Whether this mechanism could constitute a “non-cell autonomous” contributor in *C9orf72*- ALS/FTD neuropathology remains to be fully elucidated.

Overall, our study provides new insights into uptake, trafficking and release of DPR species in glia and neurons. We envision that future therapies in the context of *C9orf72*-ALS could benefit from blocking DPR uptake- and secretion-routes not only in neuronal cells but also in glia.

## Materials and Methods

### Cell culture

1321N1 astrocytoma cells were cultured in Dulbecco’s Modified Eagle Medium (Sigma) supplemented with 10% FBS (Gibco) and 5 U ml^−1^ Penstrep (Lonza). Hb9-GFP mouse stem cells were cultured as described (Wichterle *et al*, 2002) and differentiated into motor neurons with 2 μM retinoic acid (Sigma) and 1 μM Smoothened Agonist (SAG) (Millipore) for 5 days. Embryoid bodies were then dissociated with papain. All cells were maintained in a 37 °C incubator with 5% CO_2_.

### Conversion of skin fibroblasts to induced neural progenitor cells (iNPCs)

Skin fibroblasts from one healthy control (see Table 1) were reprogrammed as previously described (Meyer *et al*, 2014). Briefly, 10^4^ fibroblasts were grown in one well of a six-well plate. Day one post-seeding the cells were transduced with retroviral vectors containing Oct 3/4, Sox 2, Klf 4 and c-Myc. Following one day of recovery in fibroblast medium, DMEM (Gibco) and 10% FBS (Life Science Production) the cells were washed 1× with PBS and the culture medium was changed to Neural Progenitor Cell (NPC) conversion medium comprised of DMEM/F12 (1:1) GlutaMax (Gibco), 1% N2 (Gibco), 1% B27 (Gibco), 20 ng/ml FGF2 (Peprotech), 20 ng/ml EGF (Peprotech) and 5 ng/ml heparin (Sigma). As the cell morphology changes and cells develop a sphere-like form, they can be expanded into individual wells of a six-well plate. Once an iNPC culture is established, the media is switched to NPC proliferation media consisting of DMEM/F12 (1:1) GlutaMax, 1% N2, 1% B27, and 40 ng/ml FGF2.

Table 1. List and characteristics of control-derived cells used in this study.

| Patient sample | Ethnicity | Gender | Cell type | Age at biopsy collection |
| --- | --- | --- | --- | --- |
| 155 | Caucasian | Male | Healthy control | 40 |

### iAstrocyte differentiation and co-culture system

iAstrocytes were yielded as previously described (Hautbergue *et al*, 2017; Meyer *et al*, 2014). Briefly, iNPCs were switched to astrocyte proliferation media, which includes DMEM (Fisher Scientific), 10% FBS (Life science production) and 0.2% N2 (Gibco). Cells were grown in 10 cm dishes coated with fibronectin for 7 days unless otherwise stated. For the co-culture system, we treated iAstrocytes with 1µM DPRs for 24h, then plated Hb9-GFP mouse motor neurons on top of the astrocyte layer and kept this co-culture system for 48h before fixation and confocal imaging.

### Primary mouse cortical neurons

Primary cortical neurons were produced from E15.5 embryos of wild-type C57BL/6 mice. Brains were harvested and hemispheres were divided. In HBSS-/- medium, meninges and midbrain were removed in order to isolate the cortical tissue, which was then incubated with trypsin (Gibco) for cell dissociation. Single-cell suspension was obtained by mechanical pipetting in appropriate trituration solution (HBSS+/+ with 1% albumax, 25mg Trypsin inhibitor, 10mg/ml DNAse stock). Finally, cortical neurons were resuspended in Neurobasal medium (ThermoFisher Scientific) with B27 (Gibco), 1% Pen/Strep (ThermoFisher Scientific) and 1% Glutamine (Lonza), and seeded on Xona silicon device (#RD450) coupled with a 35mm dish previously coated with poly-D-lysine (Sigma). Cells were maintained in a 37 °C incubator with 5% CO_2_, changing medium every two days. Neurotrophic factors (2 ng/ml BDNF, 2ng/ml GDNF) were added into the medium to favour the correct directionality of axonal growth through the microgrooves. After 12 days in culture, cells were stained with Lysotracker Green (ThermoFisher Scientific) and live-imaging was performed with Airyscan microscopy (Zeiss LSM 880) at 1.5Hz for 2 minutes (561 and 488 channels; 63x 1.4NA oil immersion lens).

### Dipeptide Repeat Proteins cloning, purification and labelling

The V5-tagged 34x GA repeat was obtained using an expandable cloning strategy with Age1 and Mre1 as compatible enzymes (McIntyre *et al*., 2008). We first constructed a “start acceptor” pCi-Neo vector (Promega) by cloning a V5-3xGly/Ala insert into the Xho1/Not1 sites (ctc gag gcc acc atg ggc aaa ccg att ccg aac ccg ctg ctg ggc ctg gat agc acc ggt gca ggt gct ggc gcc ggc gga tcc gaa ttc tag ccg cgg ccg c) and a “start donor” vector with a 14xGly/Ala insert (ctc gag acc ggt gca ggt gct gga gct ggt gca ggt gct gga gca ggt gca ggt gct gga gct ggt gca ggt gct gga gca ggt gct ggc gcc ggc gga tcc gaa ttc ccg cgg ccg c) in the Xho1/Not1 sites and used these to propagate the GA repeats to construct 34 GA repeats. DNA sequences encoding V5 tag followed by 34 repeats of GA or 50 repeats of PA were subcloned in a bacterial expression vector containing an N-terminal 6xHis-Tag and a TEV protease cleavage site (pETM-11 vector, EMBL). Proteins were expressed in *E. coli* BL21 and purified on 5 mL Talon column (Clontech®) loaded with Cobalt. The proteins were eluted with a linear gradient of 12 ml from buffer A (20 mM Tris pH 7.5, 250 mN NaCl, 5mM Imidazole, 1 mM βMercaptoethanol, Glycerol 10% PMSF 0.1 mM) to buffer B (20 mM Tris pH 7.5, 250 mN NaCl, 250 mM Imidazole, 1 mM βMercaptoethanol, Glycerol 10% PMSF 0.1 mM). Eluted fractions were analyzed by SDS PAGE, and proteins were quantified spectrophotometrically using a molar extinction coefficient of ε_GA_ = 2980 M^-1^ cm^-1^ and ε_PA_ = 4470 M^-1^ cm^-1^. Proteins were assembled into fibrils at 4°C without shaking for ≤ 14 days. Proteins were labelled with Atto-550 (Atto:DPR 5:1) or Atto-647N (Atto:DPR 2:1) dyes (Atto-Tec #AD550-35, #AD647-35). Unreacted NHS-dye was removed by centrifugation (100’000*g*, 30 minutes, 4°C). Labelled assemblies were fragmented by sonication for 5 minutes in 2-ml Eppendorf tubes in a Vial Tweeter powered by an ultrasonic processor UIS250v (250 W, 2.4 kHz; Hielscher Ultrasonic, Teltow, Germany) to generate fibrillar particles that are suitable for endocytosis (with an average size of 45-55 nm), flash-frozen in liquid nitrogen and stored at -80°C. GA/PA-repeat immunoreactivity and fluorescent labelling were confirmed by resolving the generated samples by protein gel electrophoresis or dot-blot **(Fig S1A-C)**.

The morphology of DPR assemblies was assessed by Transmission Electron Microscopy in a Jeol 1400 microscope before and after fragmentation following adsorption onto carbon-coated 200 mesh grids and negative staining with 1% uranyl acetate. The images were recorded with a Gatan Orius CCD camera (Gatan). The β-sheet amyloid component of fibrillar poly-GA assemblies was assessed and confirmed by Fourier-transform infrared (FTIR) spectroscopy as described (Brasseur *et al*, 2020). Importantly, before the addition to the medium, fibrillar DPRs were sonicated for 5 min at 80% amplitude with a pulse cycle of 5 s on and 2 s off (MSE Soniprep 150); this procedure is required to disperse the aggregated β-sheet assemblies.

### Coalescence measurements

For phase-separation experiments, soluble GA and PA DPRs (stock concentration = 100 µM) were diluted in ddH2O to either 20 µM, 10 µM or 1 µM. After vortexing, the mixture was pipetted onto glass-bottom slides (Ibidi), and assembly formation was monitored over time. Images were acquired immediately, as well as after 2 and 24 hours using a Leica SP5 confocal microscope with a 63x 1.4NA oil immersion objective, 561 channel. To capture fine details for 3D rendering, assemblies were imaged with a z-stack following Nyquist sampling for optimized pixel density. Huygens Professional version 19.10 (Scientific Volume Imaging, http://svi.nl) was used to deconvolve z-stack data using the CMLE algorithm (with SNR:10 and 40 iterations) and subsequently for 3D-volume and –surface rendering thus generating **Movies S1 and S2**. The 3D rendering for **Movies S3 and S4** was performed using the software Imaris v7.7.2 (Bitplane); no deconvolution was applied here. Number, circularity and size of DPR assemblies were quantified with Fiji (Schindelin *et al*, 2012) and plotted using GraphpadPrism 8. The Fiji plug-in Trainable Weka Segmentation (Arganda-Carreras *et al*, 2017) was used to finely measure the circularity of liquid droplets in heterogeneous PA assemblies (20 µM, 24h) **(Fig S1D)**.

### Immunocytochemistry

After the addition of the DPRs, all the single-cell cultures were washed five to six times with PBS and fixed with 4% PFA for 30 min at room temperature. After fixation, cells were washed two times with PBS, permeabilized for 10 minutes with 0.1% Triton-X 100:PBS and additionally washed twice with PBS. Subsequently, cells were incubated with the blocking agent 3% BSA for 30 min and then incubated overnight at 4°C with primary antibodies (in 3% BSA). Cells were then washed 3× with PBS and incubated for 1h with the corresponding Alexa Fluor secondary antibodies (Thermo Fisher Scientific) at 1:1000 dilution (in 3% BSA) and with Hoechst when needed. Cells were washed 3x with PBS, and coverslips were mounted onto glass slides using Fluoromount aqueous mounting medium (Sigma-Aldrich). In the case of imaging with Opera Phenix® high-throughput system (PerkinElmer), coverslipping of 96-well plate is not required, and cells were imaged in PBS solution.

Primary antibodies used were: Anti-Vimentin (1:4000, Millipore Cat#AB5733), anti-αTubulin (1:1000, Sigma-Aldrich Cat#T9026), anti-CD44 (1:1000, Abcam Cat#ab157107), and anti- LAMP1 (1:25, Abcam Cat#ab25630).

### Flow cytometry

1321N1 human astrocytoma cells (1.2 x 10^6^/sample) were washed six times in PBS (to eliminate any remaining DPRs in the media), trypsinized and then resuspended in 500µl of PBS. Suspended cells were then analyzed by using an LSRII flow cytometer (BD Bioscience) with excitation at 488 nm and BD FACSDiva software (version 8.0.1; BD Bioscience) to excite the ATTO550 fluorophore. Detection of ATTO550+ signal was set at 610/20 voltage. Cells not treated with DPRs acted as control producing the gating to discriminate between the ATTO550+ and the ATTO550- cells.

### Immunoblotting

In order to collect protein extracts, iAstrocytes were washed six times in PBS (to eliminate any remaining DPRs in the media) and lysed in RIPA buffer (20 mM Tris-HCl pH 7.5, 137 mM NaCl, 10% glycerol, 1% Tryton X-100, 0.5% sodium deoxycholate, 2mM EDTA, 0.1% SDS, supplemented with protease inhibitor cocktail; Sigma-Aldrich) on ice for 20 minutes. The protein extracts were then collected in the supernatant and the concentration of each protein extract was estimated using a BCA assay (Pierce). Equal quantities of protein were mixed with 4× loading buffer (0.4 M sodium phosphate pH 7.5, 8% SDS, 40% glycerol, 10% 2- mercaptoethanol, 0.05% bromophenol blue), heated to 95 °C for 5 min, and processed for either dot-blot or western blot. Nitrocellulose membranes (0.22µm pores) were blocked in 1x TBS with 0.05% Tween (1x TBST) with 5% w/v non-fat dry milk for 1h, and then incubated with primary antibodies in 1x TBST 5% w/v non-fat dry milk at either room temperature for 2h or 4 °C overnight. Primary antibodies used were: anti-V5 (1:1000, CellSignal Cat#13202S), anti-α-tubulin (1:3000, Sigma-Aldrich Cat#T9026), anti-GA repeat (1:1000, Proteintech Cat#24492-1-AP), anti-AP repeat (1:1000, Proteintech Cat#24493-1-AP). Membranes were then washed three times for 5 min with 1x TBST and incubated with either an anti-mouse IgG-HRP-conjugate (1:5000, Bio-Rad Cat#172-1011) or an anti-rabbit IgG-HRP-conjugate (1:5000, Millipore Cat#12-348). Enhanced ChemiLuminescence (ECL) substrate was then added to the membrane to enable detection, and nonsaturated images were acquired using a G:BOX EF machine (Syngene) and Snapgene software (Syngene).

### Perturbation of endocytosis

Exposure of cells to low-temperature conditions is a commonly used method for nonspecific inhibition of endocytosis. Healthy control iAstrocytes were primed with 30 min of exposure to 4°C, and then the DPR assemblies (diluted in ice-cold DMEM) were delivered to the cells and incubated at 4°C for an additional time of 1 hour. The same procedure was applied for Alexa647-labelled transferrin, an established marker of clathrin-coated pit endocytosis (Ehrlich *et al*, 2004). In parallel conditions, cells exposed to DPR assemblies or transferrin were incubated with DMEM at 37°C for 1 hour. For the dynasore experiment, following the established protocol (Kirchhausen *et al*, 2008), cells were primed for 30 min with Dynasore (or 0.2% DMSO) and incubated at 37°C in serum-free medium before addition of transferrin or poly-GA fibrils for 1 hour. Cells were subsequently fixed in Glyoxal solution (pH=5) for 30 minutes (Richter *et al*, 2018) and imaged with confocal microscopy (Zeiss LSM 880, Airyscan mode) from the plane of sharp focus. From the images, using FIJI and creating a macro (https://github.com/paoloM1990/Quantification-of-cell-fluorescent-intensity), the *Mean grey values* of Atto550 or Alexa647 whole-cell signals were calculated. In the case of the 37-4°C endocytosis experiment, Mean grey values were log-transformed (log_10_) only for better graph visualization.

### Release of DPRs in the conditioned medium

To detect the cell release of ATTO550 DPRs into the conditioned medium, we have initially added 1µM of DPRs to the culture medium for 24h. After DPR uptake, cells were washed at least five times with PBS to remove remaining assemblies in the medium and then incubated for 24h with PhenolRed-free FBS-free DMEM. This conditioned medium (CM) was harvested in tubes which were subsequently centrifuged at 200g for 4 minutes to remove any remaining debris and dead cells. Finally, the HTS microplate reader PHERAstar FSX (BMG LABTECH) was used to measure ATTO550 fluorescence intensity (thus relative DPR concentration) in the CM on a 96-well plate. The optical module used for the fluorescence intensity measurement of each well was set to 540-20 nm (excitation light) and 590-20 nm (emission light), covering the whole area of the well. Importantly, we tested for linear dependence of fluorescence on concentration to evaluate potential inner filter effect (Fonin *et al*, 2014) by using nine known dilutions of purified DPRs in PhenolRed-free FBS-free DMEM; the plotted fluorescence values originated a standard curve with R^2^ > 0.95.

### MSD(Δt) analysis

Human iAstrocytes were exposed to 1µM ATTO550-labeled Poly-GA DPR fibrils for 24h. Cells were then washed several times with PBS and, while keeping them in 5% CO_2_ and 37°C, z-stack live-imaging was performed by taking 60 frames at ∼1frame/s rate (1 frame = 1 full z-stack) at ∼190nm lateral resolution (Zeiss LSM880, airyscan mode). Single DPR aggregates were then detected as single particles and analysed by the open-source FIJI-plug-in TrackMate (Tinevez *et al*, 2017) using difference of Gaussians (DoG) detection (*estimated blob diameter* = 0.8µm; *threshold* = 100) and Simple LAP tracker (*linking max distance* = 1µm; *gap-closing* = 1µm; *max frame gap* = 5). **Movie S5** was produced in 3D-volume rendering mode using Arivis Vision 4D software (Arivis AG, Rostock, Germany) after complete image stack deconvolution was performed for each frame with Huygens Professional version 19.10 (CMLE algorithm, SNR:20, 40 iterations). The xy data for each tracked object was analyzed to determine Mean-Squared Displacements (*MSD*) as a function of time-step, *Δt*. For untethered non-interacting objects moving freely within the media, *MSD(Δt)* is expected to evolve linearly with a gradient equal to 4D, where *D* is the Brownian translational Diffusion coefficient. If the object also experiences ballistic motion (i.e. constant translational velocity magnitude and direction), which could approximately describe microtubule transport (Van Den Heuvel *et al*, 2007), the *MSD(Δt)* becomes quadratic and is represented by the equation: *MSD = 4DΔt + v^2^Δt^2^*, where *v* is the average ballistic velocity (Dunderdale *et al*, 2012). Consequently, *MSD(Δt)* data was fitted to this expression to identify any trajectories that showed non-zero values for *v*, and so possibly reflect microtubule directed motion. For this analysis, the time range fitted was 20 seconds, and only trajectories greater than 30 seconds in length were analyzed (380 separate trajectories met this criterion).

For the *MSD(Δt)* analysis on DPR trajectories in primary mouse cortical neurons, we firstly used TrackMate for producing DPR tracks and then we implemented the MATLAB class @msdanalyzer written by Jean-Yves Tinevez (https://github.com/tinevez/msdanalyzer GitHub), already used in a previous study (Tarantino *et al*., 2014) and explained in its details (Miura & Sladoje, 2020). MSD plots and Log-Log fit plots were produced with MATLAB R2018b using the aforementioned class. A detailed explanation of how the analysis was performed (from the generation of Trackmate DPR tracks to the production of graphs after MSD analysis) can be found on Github at the following address: https://github.com/paoloM1990/Guide-to-TrackMate-Matlab-MSD

### Color deconvolution for differentiating two lysosomal populations

We developed custom scripts in MATLAB to produce color deconvolution algorithms that separated “Non-Colocalized Lysosomes” (NCLs) and “Colocalized Lysosomes” (CLs). NCLs corresponded to the signal of LysoTracker Green devoid of any overlapping with ATTO550- DPRs signal. CLs, instead, corresponded to the merged signal generated by LysoTracker Green – ATTO550-DPR colocalization. Briefly, 8-bit RGB time-lapse images were split into individual frames and a fuzzy color detection algorithm was applied to identify and isolate regions of interest. Resulting images were binarised by applying a global Otsu threshold, and then de-noised using a median filter.

### mRNA isolation and quantitative real time PCR

Total RNA from primary mouse cortical neurons (10 days in vitro) after 24h exposure to poly-GA oligomers or poly-GA fibrils (or untreated) was isolated using RNeasy Mini Kit (cat.no. 74104, Qiagen), according to the manufacturer’s manual. After reverse transcription, cDNA was processed for real time q-PCR using SYBR-green (QuantiFast SYBR Green RT-PCR Kit, Qiagen) and determination of ΔΔCT-values was done on the CFX Maestro Software (Bio-Rad Laboratories). RNA levels of GAPDH were used to standardize expression levels. For the primer sequences used in this study, see the table below:

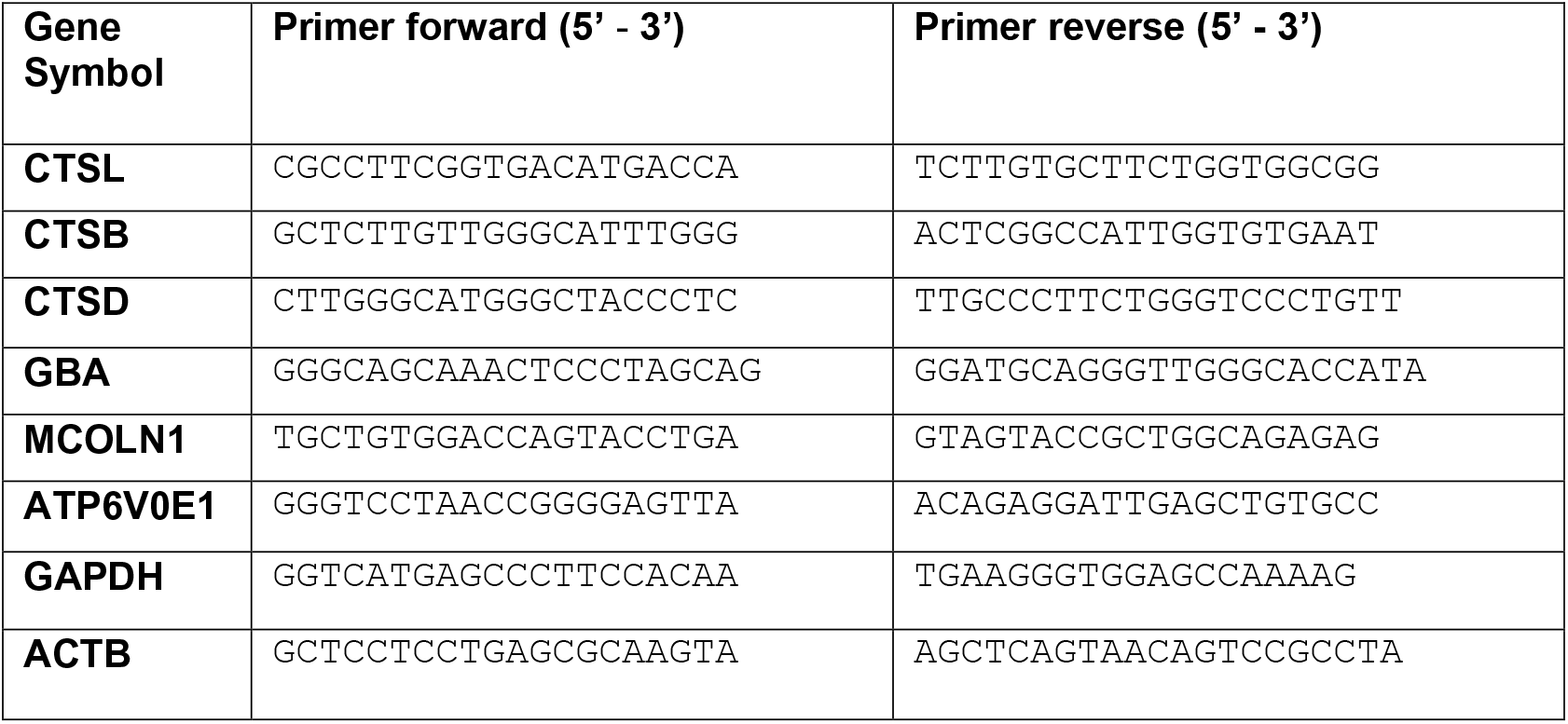

### dSTORM imaging

#### Sample preparation

High-precision (#1.5H Thickness) glass coverslips (ThorLabs, CG15CH2) were thoroughly rinsed in deionized water and dried. The glass coverslips were coated for 5 minutes at room temperature with fibronectin diluted in PBS (1:400) before iAstrocytes were plated.

#### Immunocytochemistry and staining for dSTORM imaging

Healthy control iAstrocytes were exposed to 0.5µM ATTO-647N-labeled DPR fibrils and oligomers for 24h. Cells were then washed 6 times with PBS to remove the remaining DPRs in the medium and then fixed for 60 minutes in 4% PFA (+0.2% glutaraldehyde) diluted in PBS. This long fixation period was used to minimize molecule motility (Tanaka *et al*, 2010). For dual-color dSTORM imaging, PFA-fixed cells were quenched in 50 mM NH4Cl in PBS for 5 min at RT, followed by permeabilization with 0.1% Triton-X 100 in PBS and blocking of non-specific binding sites with 2% BSA. Staining with anti-LAMP1 Mouse primary mAb (1:25, Abcam Cat#ab25630) was performed overnight at 4°C. Cells were washed with 3 x PBS for 5 minutes and incubated for 2h with Anti-Mouse IgG F(ab) ATTO488 (H+L) (HyperMOL, Cat#2112-250UG) to yield minimal linkage error (Hust *et al*, 2007). Post-fixation in 4% PFA (+0.2% glutaraldehyde) for 30 minutes was applied to further reduce molecule motility when needed. Coordinates were tracked with Nikon NIS-Element software for computational drift correction and, in a separate sample, tetraspeck beads (100 nm diameter; Invitrogen) were imaged as fiduciary landmarks for chromatic realignment. Dual-color dSTORM imaging was performed under reducing condition with Tris buffer 50mM with 10mM NaCl (pH 8), glucose (10%), cysteamine (1 M), glucose oxidase (5 mg/ml), and catalase (4 mg/ml). Approximately 8’000-10’000 frames per channel were acquired. Diffraction limited images of each channel were also acquired, providing the reference for subsequent NanoJ-SQUIRREL analysis (Culley *et al*, 2018) of image artefacts **(Fig S3C)**.

All imaging was carried out on an inverted Nikon Eclipse Ti microscope equipped with a 100× oil immersion objective (N.A. 1.49) using an Andor iXon EMCCD camera (image pixel size, 151.57 nm). ATTO-647N and ATTO-488 were imaged using 639 nm and 488 nm lasers for a 10-ms or 20-ms exposure time. We used Nikon NIS-Elements software for both image acquisition and reconstruction. After image reconstruction, the package ChriSTORM (Leterrier *et al*, 2015) was used for translating NIS-Elements localization files into compatible files for image rendering by the open-source FIJI plugin ThunderSTORM (Ovesný *et al*, 2014). Thus, ThunderSTORM enabled the ultimate visualization of data acquired by STORM imaging **(Fig S3A)**. Using the Nikon NIS-Elements software, single molecules were localised with a lateral localization accuracy of ∼20 nm for 647 channel and of ∼45 nm for 488 channel based on the Thompson equation (Thompson *et al*., 2002) **(Fig S3B)**. In addition, by using NanoJ-SQUIRREL FIJI plug-in, we have implemented block-wise FRC resolution mapping to provide local resolution measurements of our dSTORM dataset (Culley *et al*, 2018) **(Fig S3B)**.

#### Colocalization analysis in dual color dSTORM

For colocalization analysis, graphs are indicative of ∼5-6 healthy iAstrocytes with 5 ROIs taken in regions juxtaposed to the nucleus of each cell (n = 3 biological experiments). The choice of ROIs was based by excluding artefact-rich areas with the FIJI plug-in NanoJ-Squirrel (Culley *et al*, 2018).

For colocalization analysis, we used the open-source software Clus-DoC (Pageon *et al*, 2016) to generate colocalization maps (Co-Loc maps) which highlighted areas of molecular interaction between the two channels with the following parameters: L(r)-r radius = 20nm; Rmax = 500nm; Step= 10nm; Colocalization threshold = 0.4; Min colocalized points/cluster = 10.

#### Quantification and Statistical Analysis

All data are presented as means ± SEM or means ± SD, where indicated. On normally distributed data, statistical differences were analyzed using Unpaired two-tailed Student’s *t*- test (with Welch’s correction, when SDs were not equal) for pairwise comparisons or one-way ANOVA (with Tukey’s correction) for comparing groups of more than two. On non-normally distributed data, the non-parametric Kolmogorov-Smirnov test or Kruskal-Wallis test (with Dunn’s multiple comparisons) were used for pairwise comparisons or for comparing multiple groups, respectively. Normal distribution was tested with Shapiro-Wilk test and Q-Q plot. *P* < 0.05 was considered statistically significant. All graphs and tests were generated using GraphpadPrism 8.

## Acknowledgements

We thank Dr Matthew Stopford for providing iAstrocytes and Hb9-GFP motor neurons; Dr Adrian Higginbottom for providing 1321N1 human astrocytoma cells; Dr Jason King for expertise on endocytosis; Professor Elisabeth Smythe for kindly providing Transferrin647 reagent; Dr Colin Gray for support in confocal imaging and Dr Jouni Takalo for guidance on statistical analysis. Imaging work was performed at the Wolfson Light Microscopy Facility using the Inverted Nikon Eclipse Ti microscope at the University of Sheffield with the grant code MR/K015753/1. We thank the Wolfson Foundation for their support in funding the Leica Confocal microscope at SiTraN. MA is supported by the European Research Council grant (ERC Advanced Award no. 294745), MRC DPFS Award (129016), JPND-MRC (MR/V000470/1), ARUK award (ARUK-PG2018B-005) and CureAP4. LM and MA are supported by Maddi Foundation and Spastic Paraplegia Foundation. LM is further funded by a grant of the British Neuropathological Society. EFS was supported by a Motor Neurone Disease Association Prize Studentship (DeVos/Oct13/870-892 to KJDV and AJG) and Motor Neurone Disease Association project grant (DEVOS/APR18/862-79 to KJDV and AJG). KJDV was supported by the Medical Research Council (MRC) (MR/S025979/1 and MR/M013251/1 to KJDV). LB, LBr and RM were supported by the European Commission Joint Programme on Neurodegenerative Diseases (JPND-TransPathND, ANR-17-JPCD-0002-02). The present work benefited from the Electron microscopy facility of Imagerie-Gif, supported by ANR-10-INBS-04-01), and the Labex ANR-11-IDEX-0003-02. GMH acknowledges support from the Medical Research Council (MRC) New Investigator grant MR/R024162/1 and the Biotechnology and Biological Sciences Research Council (BBSRC) grant BB/S005277/1. LF is supported by the Academy of Medical Sciences (SBF002\1142) and the Medical Research Council (MRC, grant 1812144).

## Author contributions

MA, RM, PMM, LF and GMH conceived the project, designed experiments and raised the funding for the project. MA secured the PhD scholarship to support PMM project. LB, LBr and RM generated and characterized the purified DPRs. LF, MD, ACS and LMW provided iAstrocytes and Hb9-GFP motor neurons. CPW and RM performed preparation of primary mouse cortical neurons. PMM and LM performed phase separation and microfluidics experiments. CGW and DR contributed to the image analysis workflow and the interpretation of microscopy data. EFS and KJDV contributed essential reagents. SJE performed Brownian motion calculations and mathematical modelling of DPR motion. VA produced colour deconvolution algorithms for lysosomal co-localization analysis. PMM and LM wrote the manuscript, with contributions and edits from all authors.

## Supplementary Figure legends

**Figure S1.**
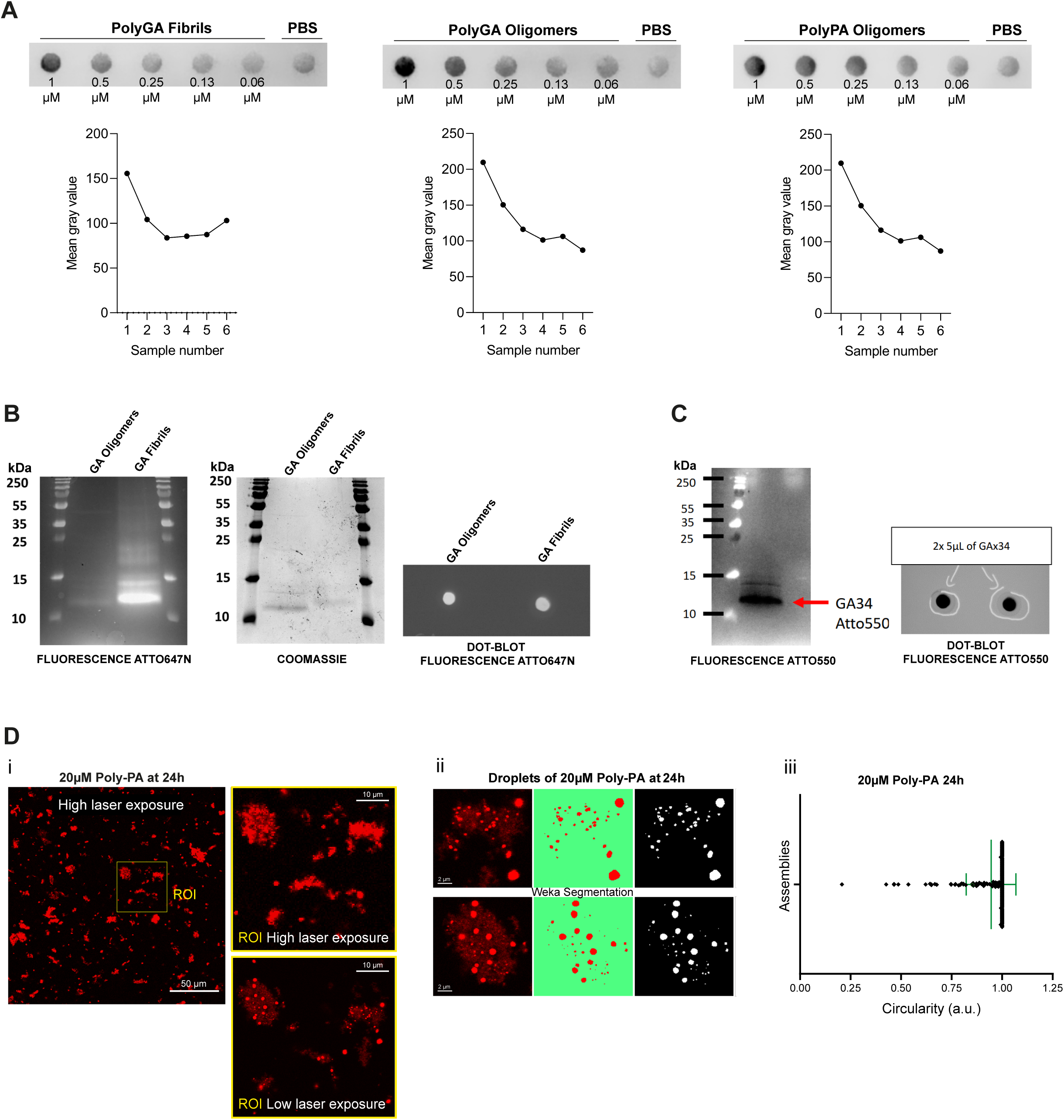
Anti-GA or anti-PA immunoreactivity and fluorophore labelling for recombinant DPRs. **A)** Dot-blotted membrane of purified poly-GA fibrils, poly-GA oligomers or poly-PA oligomers in serial dilution after staining with the respective anti-repeat antibodies (anti-GA=Proteintech 24492-1-AP; anti-PA=Proteintech 24493-1-AP). PBS indicates a control condition with no purified proteins. Graphs show quantification of Mean grey values for each corresponding condition. Data information, sample numbers: n°1 = 1µM; n°2 = 0.5µM; n°3 = 0.25µM; n°4 = 0.13µM; n°5 = 0.06µM; n°6 = PBS only. **B)** Poly-GA oligomers and fibrils labelled by ATTO- 647N (molar ratio Atto:GA = 2:1) exhibit fluorescent signal and Coomassie staining after protein gel electrophoresis and fluorescent signal after dot-blot. While both oligomers and fibrils exhibit fluorescence signal, the fibrils’ signal is three-times higher (*left*). On the coomassie gel, the quantity of monomers and fibrils is comparable (*centre*). **C)** 5 μL at 100 μM (4,5 μg) of fibrillar poly-GA labelled by ATTO-550 (molar ratio Atto:GA = 1:5) were resolved on acrylamide gel, and the fluorescence of Atto-550 (black signal) was recorded (*left*). Two samples were also analysed by dot blot on Nitrocellulose membrane, and the fluorescence of Atto-550 is displayed (black signal) (*right*). **D)** Confocal image showing phase-separated poly-PA assemblies at 20µM, 24h. The image was first acquired with high laser intensity and then with low laser intensity and Nyquist-sampling, so the droplets became distinguishable (i). After segmenting the poly-PA droplets with the Trainable Weka Segmentation FIJI plug-in (ii), their average circularity was calculated to be 0.95 (±SD) (iii); N=2.

**Figure S2.**
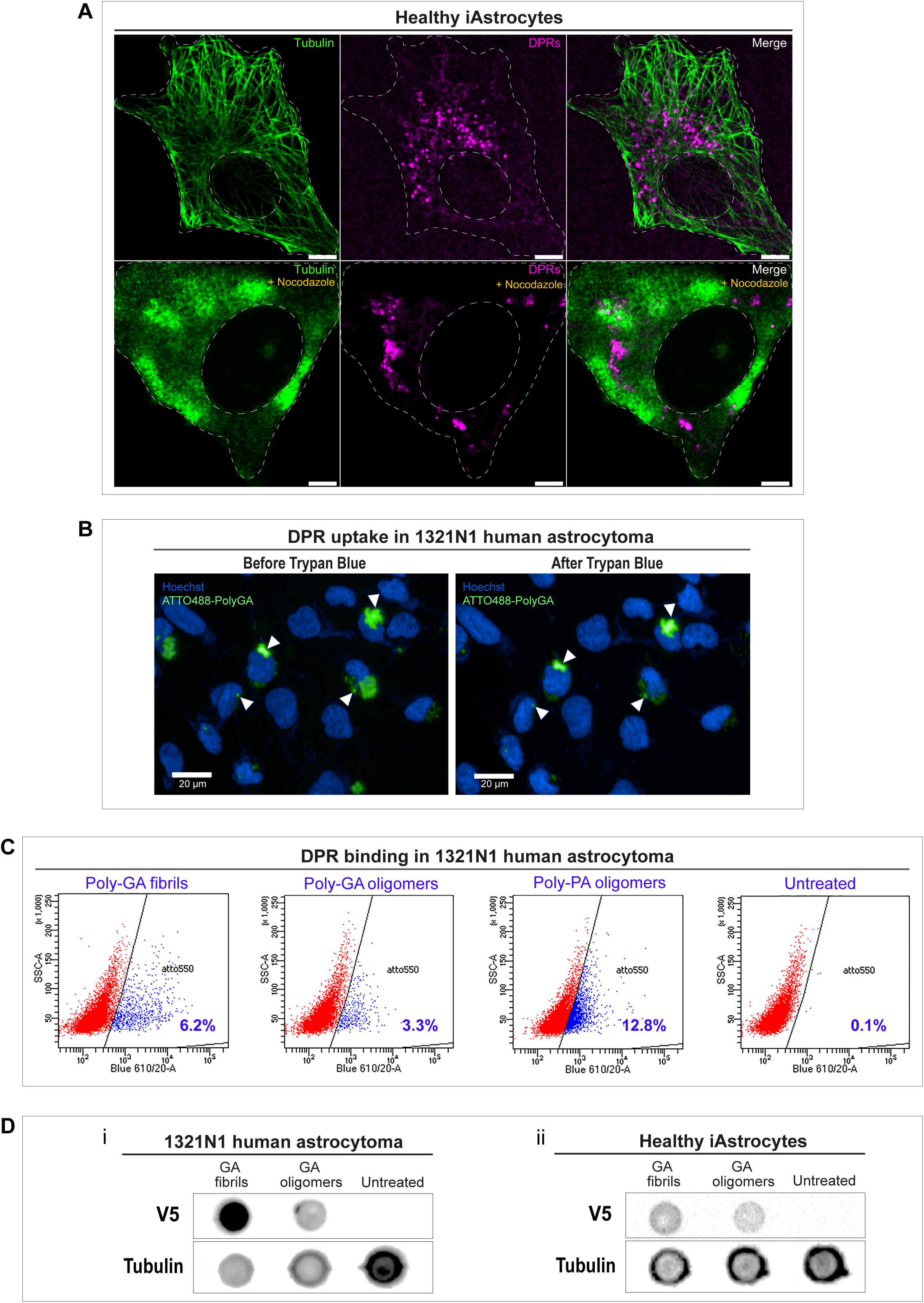
DPR uptake and binding in glia, with involvement of microtubule networks. **A)** After 24h exposure to DPRs, healthy iAstrocytes were subjected to a 30 minute pre-treatment with the microtubule de-polymerising agent nocodazole (30 µM) before fixation. Confocal images are shown, following anti-tubulin *(green)* immunostaining and ATTO550 fluorophore detection for DPRs *(magenta)*. Nocodazole treatment induces cellular relocation of DPRs. Scale bar = 10µm (upper panel), 5µm (lower panel). **B)** To confirm the uptake of Poly-GA aggregates in glia, trypan blue was used to quench any external or membrane-bound fluorescence coming from ATTO488-PolyGA added to the medium of 1321N1 human astrocytoma cells for 24h. As shown in the figure, the majority of poly-GA fluorescent signal is conserved after the addition of trypan blue (*arrow heads*), suggesting that these aggregates are taken up by the cells. **C)** Scatter plot showing ATTO550+ *(blue)* and ATTO550- *(red)* scattering events detected in 1321N1 human astrocytoma cells from a total of 10’000 events; N=3. Detection of ATTO550+ signal was set at 610/20 voltage. Cells not treated with DPRs acted as control producing the gating to discriminate between the ATTO550+ and the ATTO550- cells. **D)** After 24h DPR exposure, total protein of 1321N1 human astrocytoma cells (i) or healthy iAstrocytes (ii) was extracted into lysis buffer and then dot blotted onto a nitrocellulose membrane using a microfiltration apparatus. The membranes were then sliced into strips and analysed by anti-V5 immunostaining to show sub-populations of cells positive for V5-tagged DPRs. N=3.

**Figure S3.**
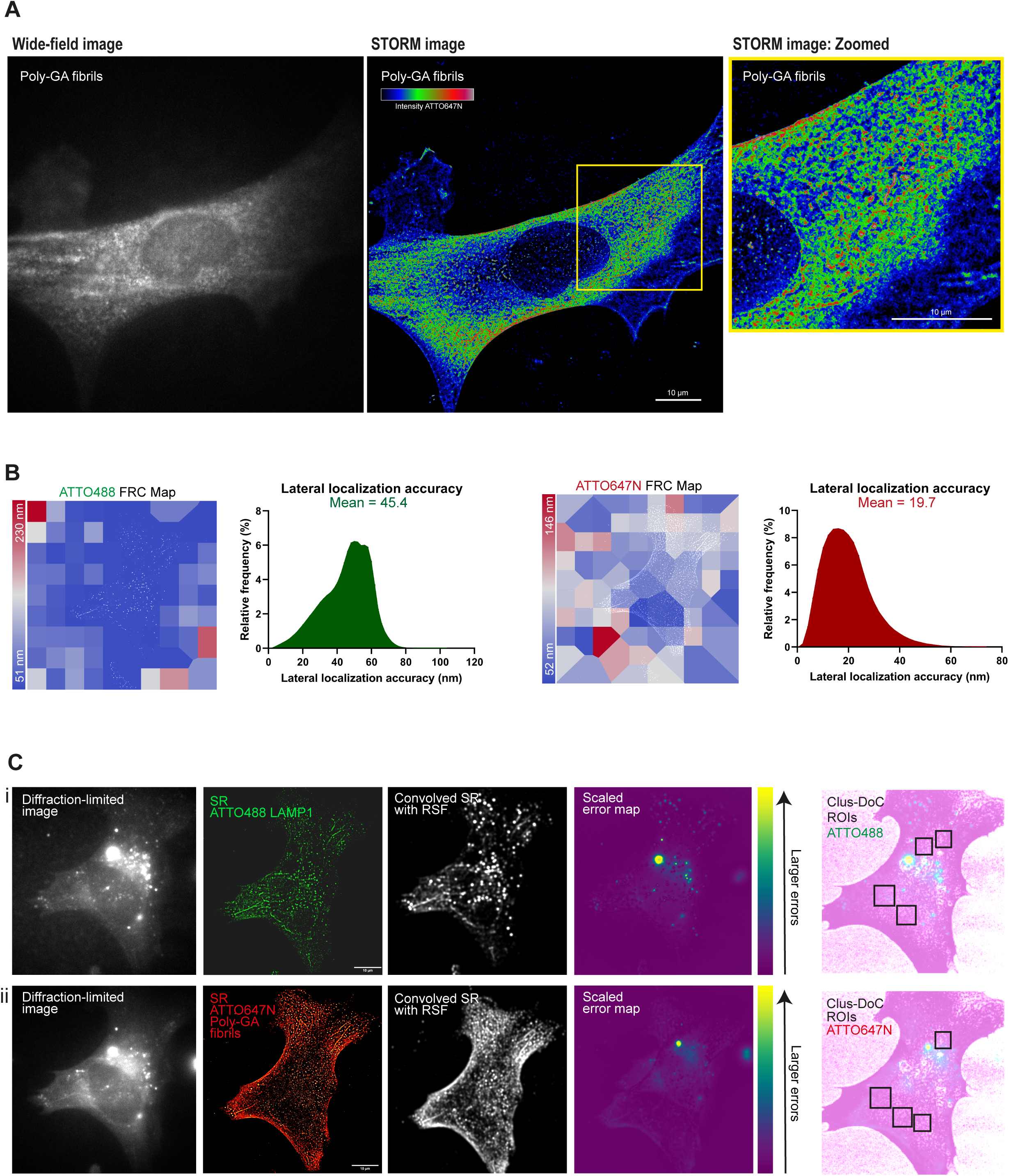
dSTORM resolution estimation and analysis of super-resolution artefacts by NanoJSquirrel. **A)** Wide-field diffraction-limited image (*left*) shows the diffuse presence of ATTO647N poly-GA fibrils across the cell with no distinguishable aggregate structures. Super-resolved image (*centre*) (8’000 frames, dSTORM) shows sub-diffractive clusters of Poly-GA fibrils with colour-coded intensity (red=maximum, blue=minimum) where major aggregate structures can now be discerned, as shown in the zoomed ROI (*right*). **B)** The lateral resolution of dSTORM images was measured implementing block-wise Fourier Ring Correlation (FRC) resolution mapping in NanoJ-SQUIRREL (Culley et al., 2018). As the resolution is anisotropic, the maps represent specific colour-coded regions of estimated resolution for each channel. The lateral resolution achieved, on average, is approximately 50-60 nm for both ATTO488 and ATTO647N channels. We also show the estimations of the lateral localization accuracy of both channels (calculated with NIS-Elements software). **C)** For lysosomes-Poly-GA fibrils co- localization analysis, diffraction-limited images of each channel were acquired. These images were used as a reference for subsequent NanoJ-SQUIRREL analysis of image artefacts in super-resolution dSTORM images. Each panel (i and ii) shows (*from left to right*): single iAstrocyte cell imaged in TIRF (reference image), super-resolution reconstruction of dSTORM data set on that same cell (’SR’), super-resolution image convolved with appropriate Resolution Scale Function (’Convolved SR with RSF’), and a quantitative map of errors between reference and convolved SR images (’Scaled error map’; colour scale indicates the magnitude of the error). Based on NanoJSquirrel-analysis of artefacts, specific ROIs for all the co-localization analysis performed in our work were chosen in artefact-free regions of the cell.

**Figure S4.**
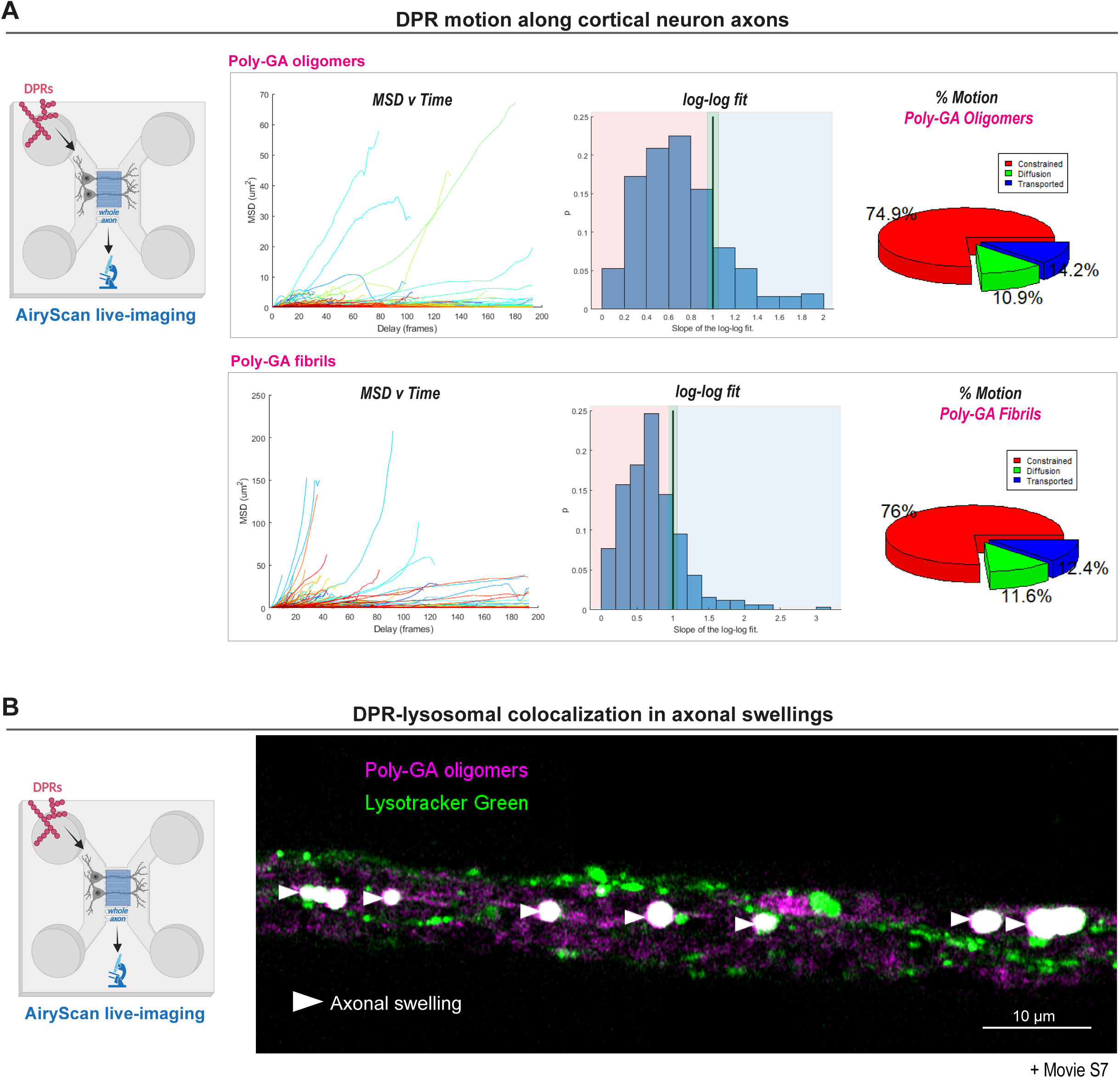
Poly-GA assemblies accumulate in axonal swellings, which are also enriched with lysosomes. **A)** Mean Square Displacement analysis of Poly-GA aggregates trajectories after live-imaging in cortical neuron axons. The plots of the MSD value as a function of time delay are shown; alongside with the plots showing the Log-Log Fit of the MSD. In a Log-Log Fit plot, the MSD curves can be approximated by straight lines of: slope 1 for diffusion motion, slope 2 for transported motion, and less than 1 for constrained motion; these are quantified and shown in the respective pie charts. ≥1000 tracks/condition; N=3. **B)** Confocal images showing accumulation of poly-GA assemblies in large axonal swellings along the axons of primary mouse cortical neurons. Upon the application of the lysosomal dye LysoTracker Green, we show that some of these axonal swellings contain Poly-GA DPRs that strongly colocalize with lysosomal organelles (*head arrows*); corresponding movie is shown in **Movie S7**.

## Supplementary Movies’ legends

### Movies S1-8

**Movie S1**

3D-volume and -surface rendering of poly-GA phase-separated oligomers at 20µM and 24h in low salt solution in the test tube. After deconvolution, 3D-surface rendering algorithms were applied to better visualize the massive and compact structure of the assemblies, along with concavity features. Deconvolution and 3D-rendering criteria are explained in the section *Method Details*_*Coalescence measurements*.

**Movie S2**

3D-volume and –surface rendering of poly-PA phase-separated oligomers at 20µM and 24h in low salt solution in the test tube. The application of 3D-rendering algorithms on deconvolved data enables to better appreciate the presence of extremely circular poly-PA droplets (average circularity = 0.95) coalescing in the test tube. The 3D-reconstructions exhibit a bulk loss of sphericity of the assemblies in the three-dimensional space concomitant with elongation along the z-axis; this is due to poor Z-axis resolution. Deconvolution and 3D-rendering criteria are explained in the section *Method Details*_*Coalescence measurements*.

**Movie S3**

3D-volume and –surface rendering of iAstrocyte-mediated uptake of poly-GA fibrils. The movie illustrates the presence of the aggregates in the perinuclear region of the cell, by showing the lateral view of the cell along the XZ axes. Red = ATTO550 poly-GA fibrils; Green = Vimentin-stained cell.

**Movie S4**

The iAstrocyte-mediated uptake of poly-GA fibrils (same of **Movie S3**) is further presented as dynamic orthogonal views (XZ, YZ) moving along the Z-stack planes.

**Movie S5**

Poly-GA fibrils undergo a mixture of Brownian motion, directed motion and constrained motion after 24h uptake in a healthy iAstrocyte cell. Poly-GA fibrils are color-coded by the signal intensity (blue=min; red=max). 60 frames were imaged at 1Hz at ∼190 nm lateral resolution in Airyscan mode. Deconvolution was applied to each frame, and the movie was generated as reported in the section *Method Details*. Fibrils were, by approximation, treated as single particles and their trajectories were analysed by Mean Square Displacement (*MSD(Δt) analysis*; **Figure 2Di**).

**Movie S6**

Accumulation of poly-GA DPRs *(magenta)* in large axonal swellings *(arrow heads)* along the axons of primary mouse cortical neurons. By zooming into few axonal swellings with higher resolution (airyscan mode), during live-imaging, we report the presence of small poly-GA proteins (by ATTO550 fluorescence) erratically moving within each axonal swelling overtime.

**Movie S7**

Poly-GA DPRs *(magenta)* and LysoTracker Green *(green)* in large axonal swellings *(arrow heads)* along the axons of primary mouse cortical neurons. The time-lapse shows that axonal swellings contain Poly-GA DPRs that strongly colocalize with lysosomal organelles.

**Movie S8**

The movie shows a three-dimensional view of co-localization between all the DPRs *(magenta)* and the lysosomes *(green)* contained in the proximal axons of mouse cortical neurons. Each region-of-interest (ROI) is delimited by a white square. On the right panel of the movie, each ROI is presented in a zoomed view moving through the Z-planes of the Z-stack dataset. This shows that co-localization between DPRs and lysosomes is consistent throughout the three-dimensional space.

